# How Connecting the Legs with a Spring Improves Human Running Economy

**DOI:** 10.1101/2023.04.03.535498

**Authors:** Jon P. Stingel, Jennifer L. Hicks, Scott D. Uhlrich, Scott L. Delp

## Abstract

Connecting the legs with a spring attached to the shoelaces reduces the energy cost of running, but how the spring reduces the energy burden of individual muscles remains unknown. We generated muscle-driven simulations of seven individuals running with and without the spring to discern whether savings occurred during the stance phase or the swing phase, and to identify which muscles contributed to energy savings. We computed differences in muscle-level energy consumption, muscle activations, and changes in muscle-fiber velocity and force between running with and without the spring. Across participants, running with the spring reduced the measured rate of energy expenditure by 0.9 W/kg (8.3%). Simulations predicted a 1.4 W/kg (12.0%) reduction in the average rate of energy expenditure and correctly identified that the spring reduced rates of energy expenditure for all participants. Simulations showed most of the savings occurred during stance (1.5 W/kg), though the rate of energy expenditure was also reduced during swing (0.3 W/kg). The energetic savings were distributed across the quadriceps, hip flexor, hip abductor, hamstring, hip adductor, and hip extensor muscle groups, whereas no changes in the rate of energy expenditure were observed in the plantarflexor or dorsiflexor muscles. Energetic savings were facilitated by reductions in the rate of mechanical work performed by muscles and their estimated rate of heat production. The simulations provide insight into muscle-level changes that occur when utilizing an assistive device and the mechanisms by which a spring connecting the legs improves running economy.

## I. INTRODUCTION

**M**ILLIONS of people run for exercise, transportation, or leisure. Humans adopt energetically efficient movement strategies, especially in repetitive tasks such as running [1]–[3], enabling them to run long distances while expending a surprisingly small amount of energy [4], [5]. Storing and releasing elastic energy in our muscles and tendons is an energy-saving mechanism [5]–[7]. This elastic energy release enables our muscles to more efficiently support body weight and modulate center-of-mass velocity; however, vertical support and forward propulsion still constitute up to 80% of the net energy expended in running [8], [9].

Several studies have used elastic elements to influence runners. Using a damped spring model of running, McMahon et al. [10] discovered that changing the compliance of the track under a runner’s feet could increase running speed while also decreasing measures associated with injury. A shoe capable of returning more elastic energy from ground impact reduces the energetic cost of running by 4% [11]. Conversely, highly cushioned shoes increase leg stiffness and impact loading on the runner [12]. Each of these approaches focuses on the interaction of the runner with the ground.

Simpson and colleagues [13] designed an “exotendon”—a spring connecting the shoelaces—that improved running economy by storing and releasing energy primarily during the flight phase. The exotendon reduced energy expenditure by 6.4% when running at a constant speed [13]. When using the device, runners increased their stride frequency and reduced their biological hip flexion, hip extension, and knee extension moments [13]. The reductions in hip flexion and extension moment peaks occurred when the exotendon applied its maximum force, which resulted in device-generated hip extension and flexion moments on the leading and trailing limbs, respectively. Simpson et al. [13] postulated that by assisting the acceleration and deceleration of the limb during the swing phase, the exotendon enabled a faster stride frequency, which reduced the required knee extension moment during stance. Despite the changes in hip and knee moments, no clear patterns emerged in how measured muscle activity changed, and the muscle-level explanation for energy savings remains unclear.

Simulations can estimate muscle dynamics during movement, which can elucidate how muscle energetics change when using a device. Specifically, simulations allow us to explore changes in muscle fiber length, velocity, and force, which are challenging to measure experimentally [14]–[16]. For example, simulations have provided insights on how spasticity affects the response of muscles during stretches and in gait [17], plantarflexor muscle-tendon dynamics and energetics for rearfoot and forefoot striking during running [18], and the effects of elastic ankle exoskeletons [19]. Further, they have been used to better understand the energetics of walking on level ground, on an incline, with load, and with assistive devices [20]–[23].

The purpose of this study was to investigate muscle-level changes that enable humans to save energy when running with an exotendon connecting their legs. Based on the decreases in muscle-generated joint moments observed by Simpson et al. [13], we hypothesized that while running with the exotendon, users save energy across both the stance and swing phases of gait due to the reduced joint moments, and that the savings occur in the hip flexor, hamstring, and quadriceps muscle groups.

## II. Materials and Methods

### A. Participants

Seven healthy young adults, without musculoskeletal or cardiopulmonary impairments, participated in the study (4 female, 3 males; age: 25.7±1.1 years; height: 171.9±7.8 cm; mass: 63.4±7.0 kg; mean ± standard deviation). Participants were excluded if they did not run at least once per week or if they had experienced a lower-extremity injury in the preceding six months. The study was approved by the Stanford University Institutional Review Board, and all participants provided written informed consent prior to participation.

### B. Experimental protocol

Participants ran with the exotendon during three visits to a motion capture laboratory: two acclimation visits followed by an evaluation visit. On the first visit, we fabricated an exotendon customized for the participant’s leg length [13]. During the first and second visits, participants underwent a structured training protocol to learn how to use the exotendon. Participants first acclimated to the device by completing at least four 15 m bouts of both walking and running. Once the participants were comfortable, they were fitted with breath-by-breath indirect calorimetry equipment (Quark CPET, COSMED, Rome, Italy) that measured the whole-body rate of energy expenditure for the remainder of the training session. Participants were instructed to consume only water in the 3 hours prior to the data collection [24]. They completed a five-minute static standing trial to measure the baseline rate of energy expenditure. Next, participants completed four randomized, alternating 10-minute runs with and without the exotendon, with five-minute rests between. We refer to runs without the device as ‘natural’ running and those with the device as ‘exotendon’ running. The speed of each run was 2.67 ms^-1^; this speed was chosen to match previous experiments [13]. All runs were completed on an instrumented split-belt treadmill (Bertec Corporation, Columbus, OH, USA).

Only participants who saved at least a 3% average rate of energy expenditure during exotendon runs compared to their natural runs by the second visit proceeded to the third visit (two participants did not meet this threshold). This threshold helped ensure those participants completing the full protocol would save energy during the final data collection. The third visit occurred after up to two rest days following the second training session. On this day, we collected kinematic, kinetic, and electromyographic (EMG) data in addition to the indirect calorimetry data for all runs. We collected EMG signals for 13 lower limb muscles (Trigno IM, Delsys Inc., Natick, MA, USA) at 2000 Hz, including the adductor group, rectus femoris, vastus lateralis, vastus medialis, gluteus maximus, gluteus medius, medial and lateral hamstrings, tibialis anterior, medial and lateral gastrocnemius, soleus, and peroneus muscles. Participants performed a series of maximum-effort trials to calibrate the EMG signals, including five maximum height jumps, five sprints, and a series of isometric and isokinetic contractions for the hamstrings, hip abductors, hip adductors, hip flexors, tibialis anterior, and peroneus [25]. We bandpass filtered EMG signals at 30-500 Hz (4th order, zero-phase shift Butterworth), rectified, and low-pass filtered them at 6 Hz (4th order, zero-phase shift Butterworth). Signals were then normalized by the highest value recorded for a given muscle in the maximum activation trials.

We placed 40 retro-reflective markers on participants for 3D motion capture, which was recorded at 100 Hz using an 11-camera optical motion capture system (Motion Analysis Corporation, Santa Rosa, CA, USA). Anatomical markers (22) were placed bilaterally on the second and fifth metatarsals, calcanei, malleoli, femoral epicondyles, anterior and posterior superior iliac spines, acromia, C7 vertebrae, and sternum. Tracking markers (16) were placed on the shanks and thighs of the legs. Two markers were placed on the shoes where the exotendon was connected. Markers on the medial femoral epicondyles and malleoli were removed after a static standing trial. Participants performed a hip circumduction exercise to estimate the location of the hip joint center [26]. On this evaluation day, participants performed four seven-minute runs with five-minute breaks, alternating conditions between natural and exotendon following the same randomized order as their previous two visits.

### C. Musculoskeletal modeling

We adapted a generic musculoskeletal model in OpenSim 4.3 with 37 degrees of freedom [16], [27], [28]. We removed the arms and associated degrees of freedom, and we locked the metatarsophalangeal joints, leaving 21 degrees of freedom. The generic model was scaled to match each participant’s anthropometry using a standing static trial.

The model incorporated 80 Hill-type muscle-tendon units to actuate the lower extremities [28]–[30]; each muscle’s maximum isometric force was adjusted by scaling the participant’s total muscle volume based on mass and height [31]. The exotendon was modeled as a linear path spring with a customized slack length (measured from each exotendon) and stiffness (characterized from the material) for each participant [13]. The path spring connected the calcanei at the location measured during motion capture.

For each participant and condition, we examined four gait cycles. We low-pass filtered (15 Hz, 4th order, zero–phase shift Butterworth) the motion capture and ground reaction force and moment data. For each gait cycle, we estimated 3D kinematics using the Inverse Kinematics tool in OpenSim [16]. These kinematics; ground reaction forces and moments; and scaled musculoskeletal model were used as inputs to OpenSim’s Residual Reduction Algorithm (6 Hz low-pass filtered), which modified the location of the torso center of mass and provided body segment mass adjustments to reduce dynamic inconsistencies [14]. We ran the algorithm on the eight gait cycles for each participant and averaged all adjustments [22]. These adjusted models were used for all subsequent simulations of each gait cycle.

### D. Muscle-driven simulations

We used the MocoTrack tool in OpenSim Moco, which utilizes optimal control methods, to generate dynamic, muscle-driven simulations of each gait cycle [32]. In each simulation we provided the kinematic trajectories, ground reaction forces and moments, and adjusted musculoskeletal model as inputs. Ground reaction forces and moments were prescribed, and the optimizer weightings encouraged close tracking of input kinematics, while allowing for slight deviations due to inconsistencies between the musculoskeletal model, ground reaction forces, and kinematic data. The primary objective in the optimization was to minimize the deviation from the input kinematic trajectories in addition to the sum of squared muscle excitations. Reserve actuators were added to each of the joint coordinates in the model but were heavily penalized in the optimization cost function to ensure that the muscles were primarily generating the motion. We included an additional secondary cost term to penalize taking advantage of an initial state with high muscle activations. The final simulation is a dynamically consistent set of kinematic trajectories and muscle states (i.e., activations, lengths, etc.). The model and simulation code are publicly available at (https://simtk.org/projects/exotendon_sims).

### E. Model of energy expenditure

We measured the average rate of gross energy expenditure using indirect calorimetry during the final two minutes of each running or static standing trial [33]. We subtracted the baseline rate of energy expenditure, measured during static standing, from the running trials to obtain the net whole-body rate of energy expenditure for each condition and run. We then computed the average percent change in the net whole-body rate of energy expenditure from the natural running to the exotendon running conditions across the 7 participants.

To examine the rate of energy expenditure in each muscle we utilized a model previously described by Uchida et al. [34], [35]. This model estimates a muscle’s rate of energy expenditure based on the muscle fiber length, velocity, force, and activation and was used to estimate the energy expenditure for 40 muscles in the lower limb. We computed values for a single limb and assumed symmetry between limbs. The simulated whole-body rate of energy expenditure was computed by integrating the sum of the individual muscle rates of energy expenditure over the full gait cycle, dividing by the duration of motion, and multiplying by two for the second limb. We averaged the energy rate across the four simulated gait cycles for each subject and condition.

We combined functional muscle groups to provide a single estimate of energy expenditure for the hip flexor, hip adductor, hip abductor, hip flexor, quadriceps, hamstrings, dorsiflexor, and plantarflexor muscle groups. We computed the average rate of energy expenditure for each muscle group during both natural and exotendon running. We confirmed a normal distribution using a Shapiro-Wilk test and used a paired two-sample t-test to determine differences in each group between conditions (α=0.05). We used the simulated whole-body rate of energy expenditure from the natural condition to compute percentage changes between conditions.

To compute the average total muscle metabolic rate for the stance phase, the sum of all the individual muscles’ rate of energy expenditure in one leg was integrated over the time when the foot was in contact with the ground, divided by the duration of contact time. Average total muscle metabolic rate for the swing phase was similarly computed over the time when the foot was not in contact with the ground. We computed the averages for natural and exotendon running in both stance and swing, and confirmed the data had a normal distribution with a Shapiro-Wilk test. We used a paired two-sample t-test to determine differences between stance and swing values in each condition (α=0.05). For all percent-change calculations, we used the participants’ average simulated whole-body natural rate of energy expenditure to scale the result.

### F. Simulation verification

#### a. Simulated vs. measured metabolic rate

The RMS and peak error in the average net whole-body metabolic rate across all 14 of the subject-conditions were 1.13 W/kg and 2.50 W/kg, respectively (Fig. 1). The measured rate of energy expenditure was 0.9 W/kg (8.3%) lower during the exotendon trials compared to the natural condition. It was 1.4 W/kg (12.0%) lower in the simulations, and the simulations correctly detected the reduction in energy expenditure in all seven participants.

**Fig. 1.**
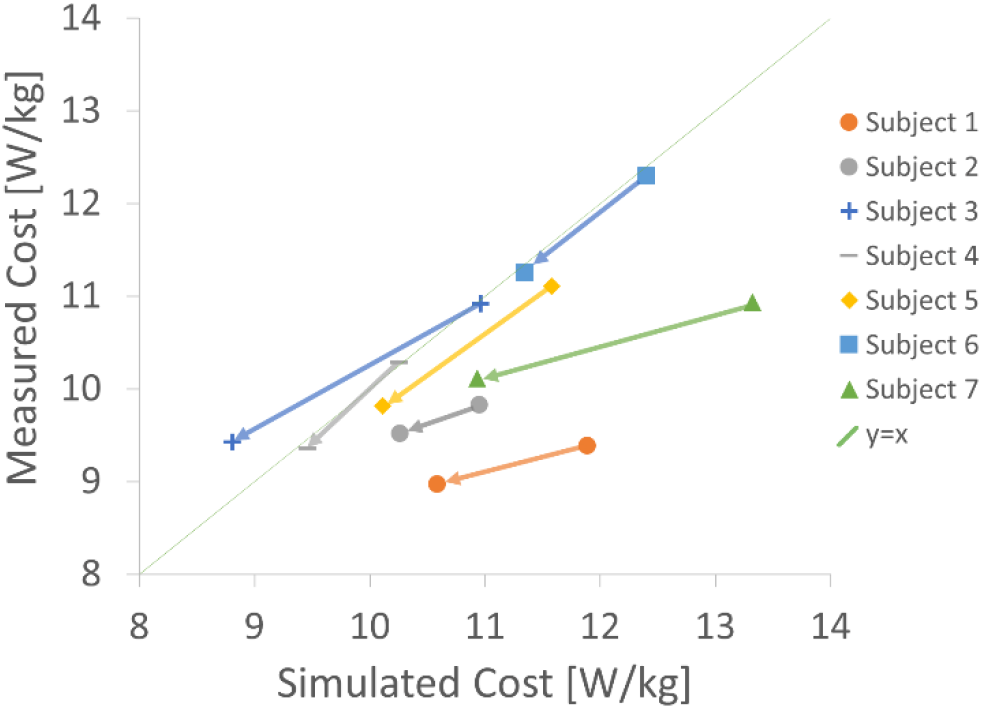
Simulated vs. measured net whole-body rate of energy expenditure for each participant. The arrows for each participant designate the change from natural to exotendon runs. A perfect estimate of the simulated cost would fall along the y=x line. Our simulations correctly identified that the exotendon reduced energy cost for all participants.

#### b. Kinematics and kinetics

The root-mean-square (RMS) coordinate angle difference between simulated and inverse kinematics motions averaged across participants, gait cycles, and coordinates was 1.2°; the maximum difference at any time was 10.5° in the lumbar extension angle. The RMS difference in the pelvis coordinate locations was 1.3 cm, with a maximum difference at any time of 7.5 cm. The peak and RMS reserve moments across all simulations were 6.26 and 0.07 Nm (6.5% and 0.1% of the peak net joint moment), respectively, which are near the 5% threshold recommended in Hicks et al. [36]. The peak and RMS residual forces across all simulations were 0.34 and 0.04 N (each less than 0.1% of peak applied net ground reaction force), respectively, which falls below the recommended 5% [36]. The largest peak and RMS residual moment across all simulations was 5.10 and 0.44 Nm (0.4% and 0.03% of the product of the center of mass height and the applied net ground reaction force), respectively, which falls below the recommended 1% [36].

#### c. Muscle activations

We qualitatively compared the simulated muscle activations to measured EMG for major muscle groups including the hip extensors, hip adductors, hip abductors, hamstrings, quadriceps, plantarflexors, and dorsiflexors (Fig. 2). We compared the simulation to EMG using a weighted average of the muscles within each group, using the maximum isometric muscle force as the weight. The simulations capture salient features of the timing of muscle activation, as well as trends in how activation changed when running with the exotendon (e.g., decreased quadriceps activity with the exotendon). Simulated muscle activations tended to be larger than EMG signals seen in the experimental data, likely because the simulated muscles have less capacity to generate force than the participants’ muscles [37]. We used EMG for qualitative validation and did not statistically test for between-condition differences.

**Fig. 2.**
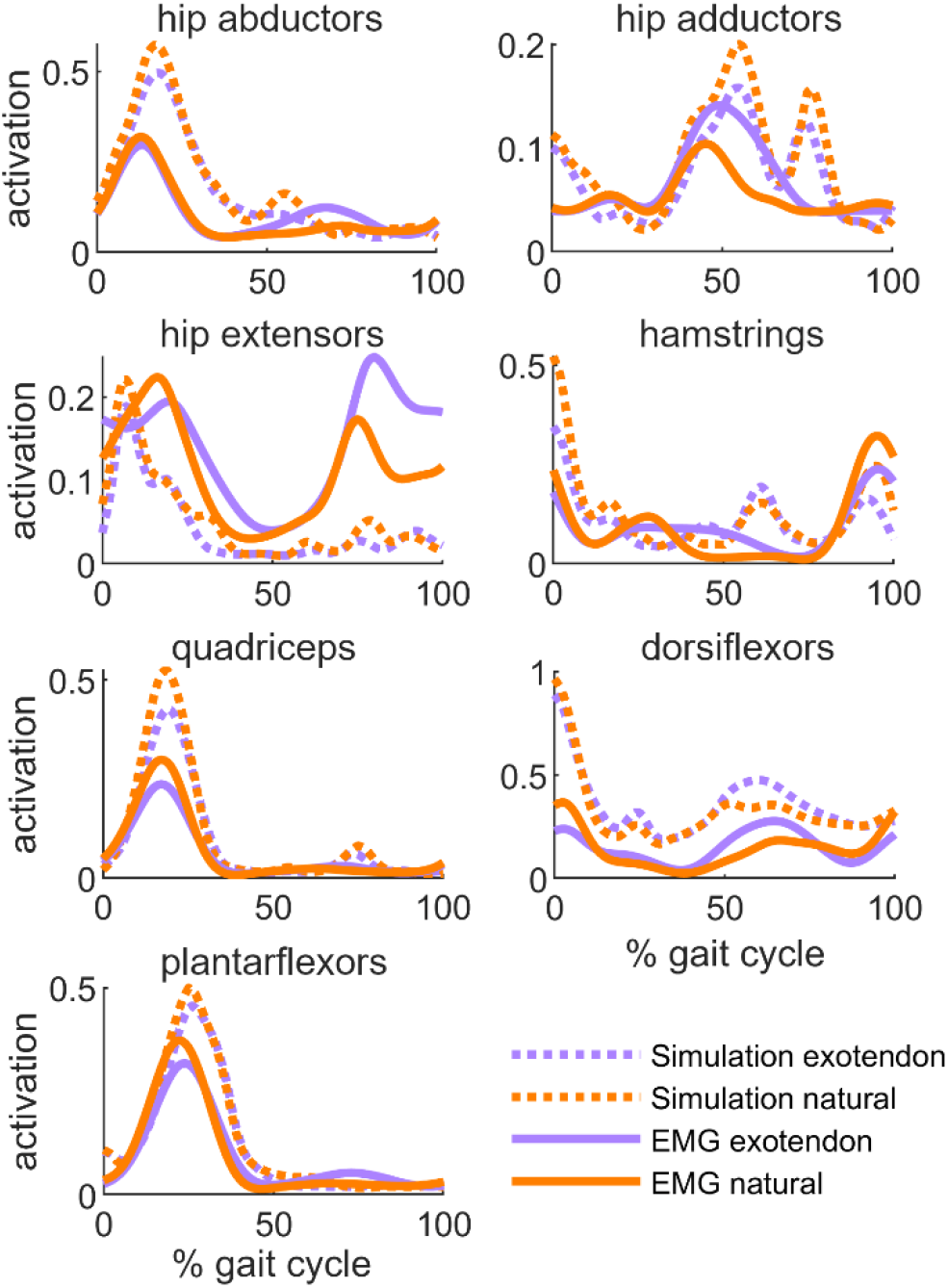
Simulated and measured muscle activity is shown for natural (orange) and exotendon (purple). Simulated activation and EMG signals were averaged across all participants. EMG signals are normalized by the highest signal recorded for a given muscle in the maximum activation trials. Note there were no EMG signals for the hip flexors.

#### d. Sensitivity analysis

We performed a sensitivity analysis of the convergence tolerance used in our optimizations to confirm that a stricter tolerance on the optimization would not change the results. When tightening the convergence tolerance from 0.01 to 0.001, the objective values did not change more than 0.3%. This small change indicated that the value of 0.01 was suitable for the problem, and that tightening the tolerance would likely not significantly change the result.

## III. Results

Participants running with the exotendon reduced their measured rate of energy expenditure by an average of 0.9±0.2 W/kg (8.3±1.3% mean ± standard error, *P*=.001). In simulation, we computed a 1.4±0.2 W/kg (12.0±1.8%, *P*=.001) reduction in energy expenditure rate on average. When comparing exotendon to natural, runners spent 1.5±0.2 W/kg (12.8±1.2%, *P*<.001) less in stance and 0.3±0.1 W/kg (2.5±0.9%, *P*=.04) less in swing (Fig. 3).

**Fig. 3.**
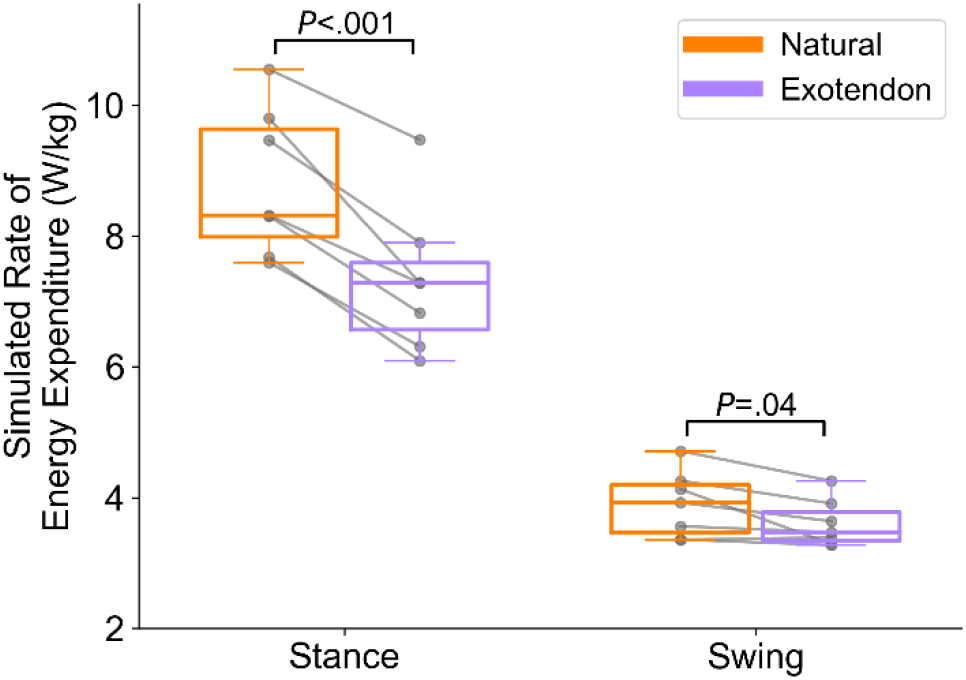
Stance- and swing-phase simulated rate of energy expenditure for natural (orange) and exotendon running (purple). A line connects each subject’s data with and without the exotendon.

The quadriceps muscles saved the most energy on average, contributing a reduction of 0.14±0.04 W/kg (*P*=.02), or 1.2±0.4% per leg as a percentage of the whole-body metabolic rate in the natural condition (Fig. 4). Additionally, the hip flexors (0.11±0.02 W/kg, 1.0±0.2%, *P*=.002), hip abductors (0.1±0.02 W/kg, 0.8±0.2%, *P*=.004), hamstrings (0.09±0.03 W/kg, 0.8±0.3%, *P*=.03), hip adductors (0.06±0.01 W/kg, 0.6±0.1%, *P*=.001), and hip extensors (0.05±0.02 W/kg, 0.4±0.1%, *P*=.046) reduced their average rate of energy expenditure. Only the plantarflexor and dorsiflexor muscles did not significantly change between conditions.

**Fig. 4.**
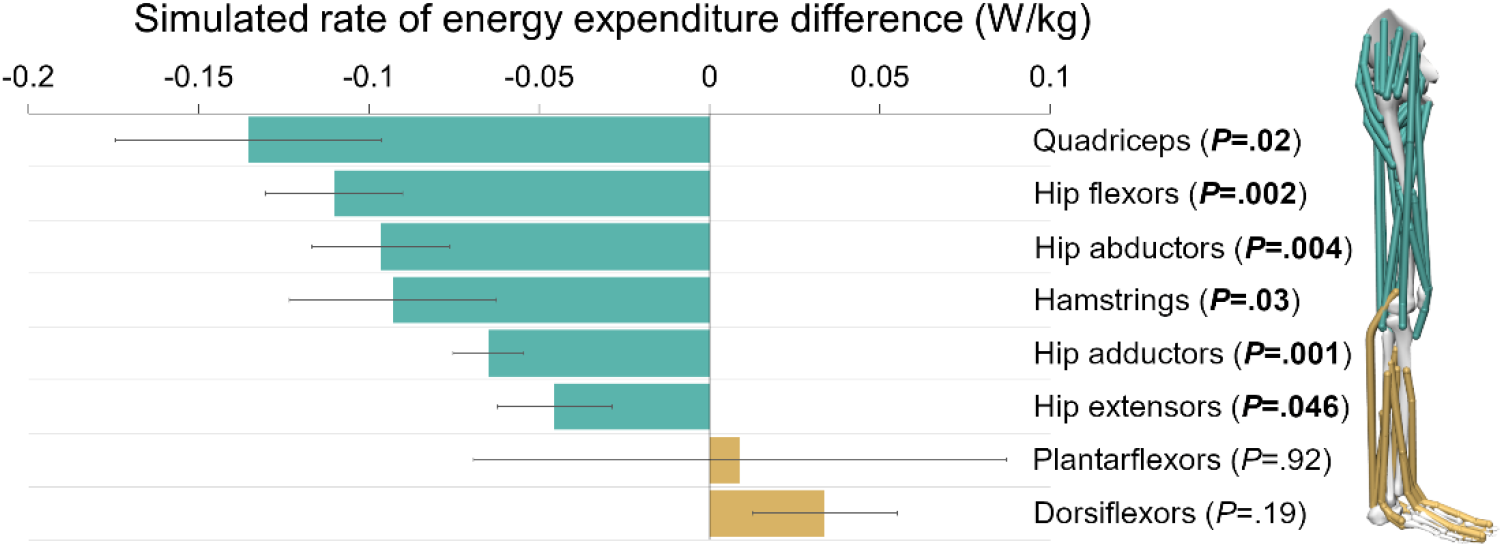
Changes in simulated energy expenditure from natural to exotendon runs for major muscle groups (bar: mean, error bar: standard error). The values shown are for one leg. A negative change (green) denotes energy savings for the muscle group in exotendon runs, and a positive change (gold) denotes more energy being spent.

We examined individual muscle metabolic rates, activations, fiber velocities, and active fiber forces for the top three muscle groups that saved energy—the quadriceps, hip flexor, and hip abductor muscles—to better understand the mechanisms of savings. Further, we evaluated the range of motion and peak moments of the degrees of freedom actuated by these muscles to explore the joint-level kinematic and kinetic changes associated with the muscle-level energetic savings. We did not perform any statistical analysis on the individual simulated muscle energy expenditures, activations, fiber velocities, active fiber forces or the joint-level kinematics and kinetics; these are exploratory analyses of the changes that may influence the statistically tested changes in metabolic rate.

Within the quadriceps muscle group, the vastus lateralis saved the most energy, with unilateral savings of 0.07±0.03 W/kg (0.7±0.3%, *P*=.04). The vastus lateralis expended energy primarily during stance. Peak muscle activation, peak fiber velocity, and peak active fiber force were all reduced during exotendon running (Fig. 5). The peak knee extension moment decreased from natural to exotendon by 0.40±0.13 Nm/kg on average. The knee flexion-extension range of motion decreased with the exotendon from 78±3° (natural) to 63±3° (exotendon) on average.

**Fig. 5.**
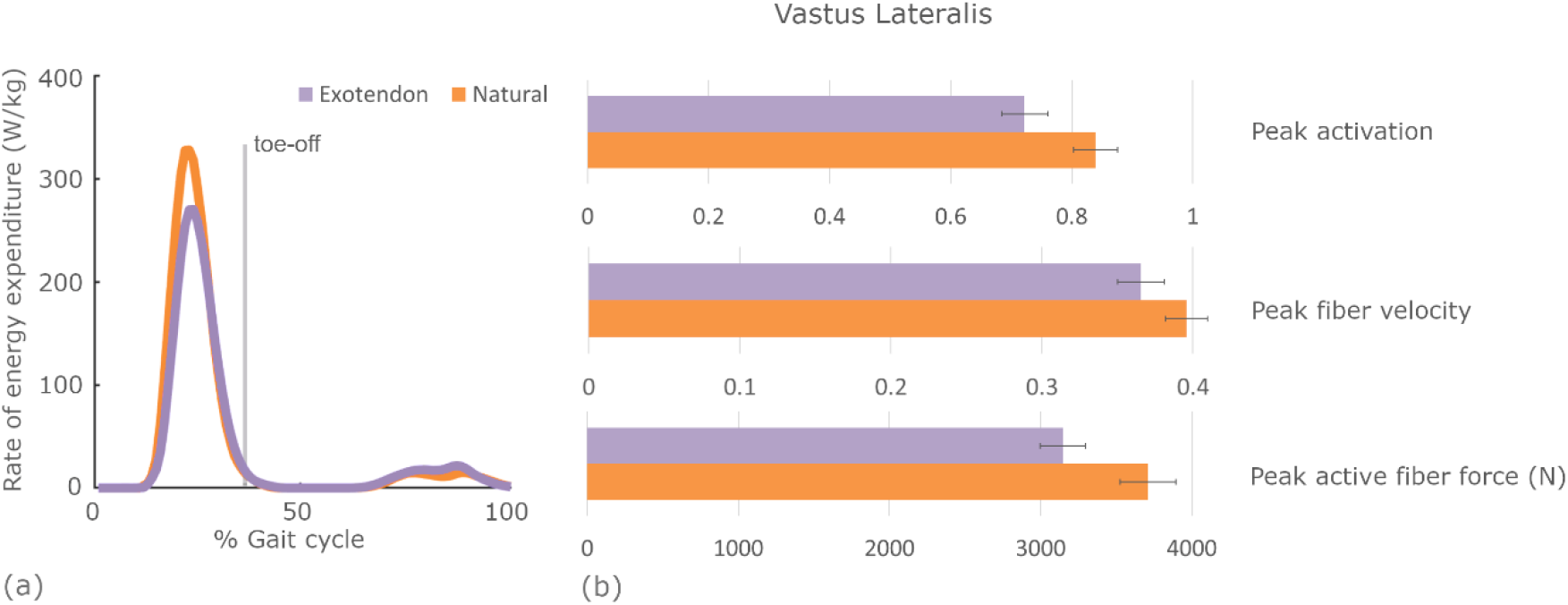
Changes in the rate of energy expenditure in the vastus lateralis throughout the gait cycle and the corresponding muscle state changes that contribute to the reduction. (a) The average vastus lateralis muscle rate of energy expenditure is plotted throughout the gait cycle for both natural (orange) and exotendon (purple) runs. (b) The peak muscle activation (ranging from 0–1), fiber velocity (fiber lengths per second), and active fiber force averaged across all participants at the time of highest energy expenditure for both natural (orange) and exotendon (purple) running.

Within the hip flexor muscle group, the psoas saved the most energy, with unilateral savings of 0.05±0.01 W/kg (0.5±0.2%, *P*=.007). Unlike the quadriceps, the psoas primarily expended energy during the swing phase. We found that it reduced peak muscle activation, peak fiber velocity, and peak active fiber force during exotendon running (Fig. 6). The peak hip flexion moment decreased from natural to exotendon by 0.25±0.04 Nm/kg on average. The hip flexion-extension range of motion decreased from 50±2° (natural) to 44±1° (exotendon) on average.

**Fig. 6.**
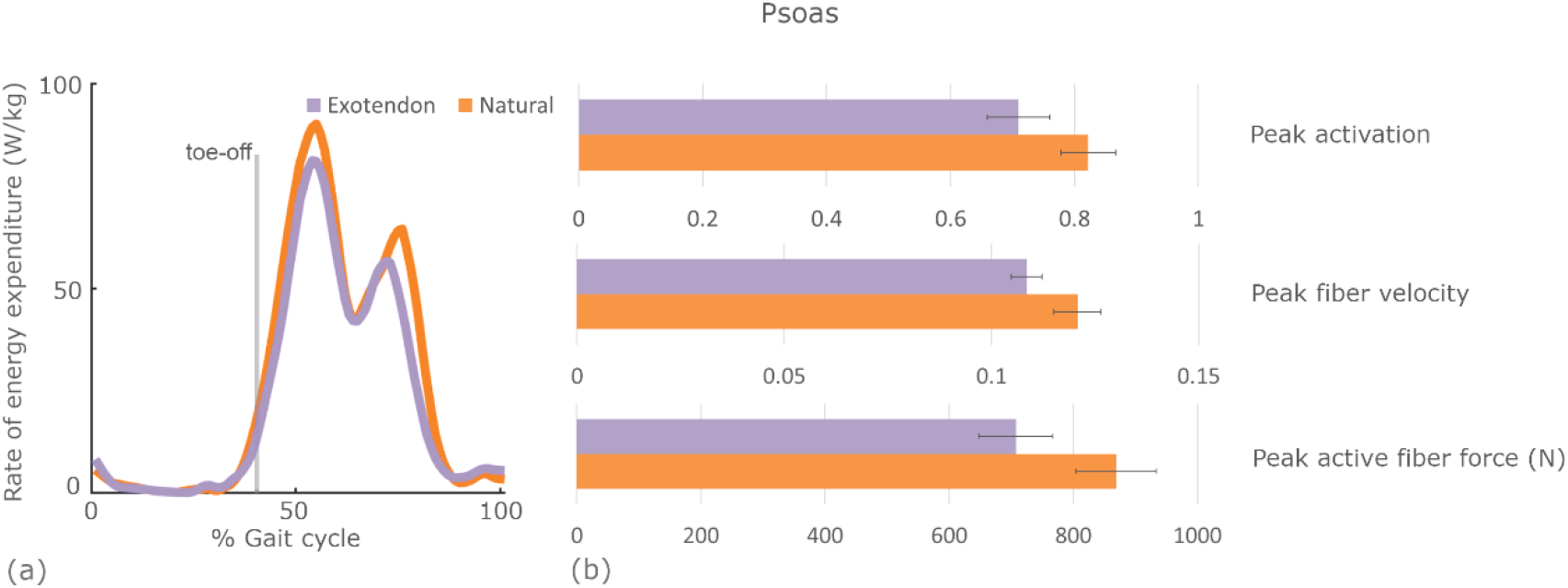
Changes in the rate of energy expenditure in the psoas throughout the gait cycle and the corresponding muscle state changes that contribute to the reduction. (a) The average psoas muscle rate of energy expenditure is plotted throughout the gait cycle for both natural (orange) and exotendon (purple) runs. (b) The peak muscle activation (ranging from 0–1), fiber velocity (fiber lengths per second), and active fiber force averaged across all participants at the time of highest energy expenditure for both natural (orange) and exotendon (purple) running.

Within the hip abductor muscle group, the gluteus medius saved the most energy, with unilateral savings of 0.06±0.01 W/kg (0.5±0.1%, *P*=.008) of the full-body rate. The gluteus medius primarily expended energy during stance. Peak muscle activation, peak fiber velocity, and peak active fiber force were all reduced during exotendon running (Fig. 7). The peak hip abduction moment decreased from natural to exotendon by 0.15±0.05 Nm/kg on average. The hip adduction-abduction range of motion decreased from 22±1° (natural) to 17±1° (exotendon) on average.

**Fig. 7.**
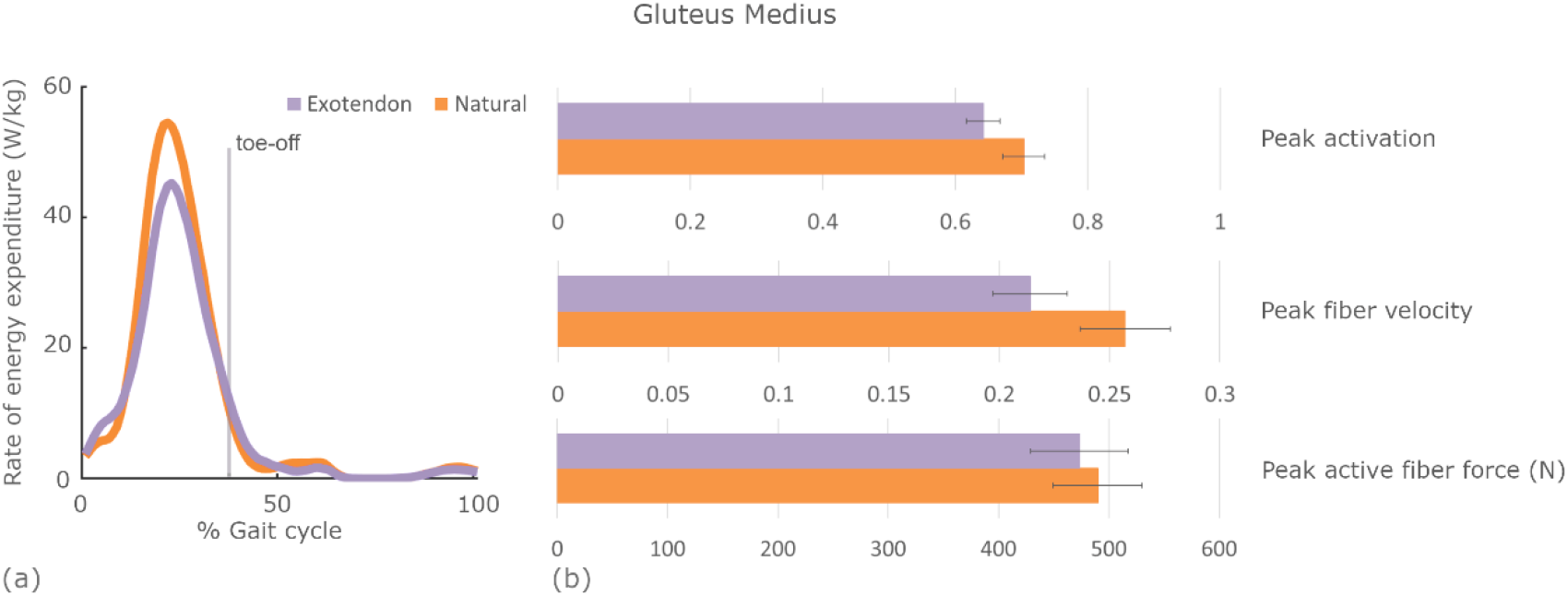
Changes in the rate of energy expenditure in the gluteus medius throughout the gait cycle and the corresponding muscle state changes that contribute to the reduction. (a) The average gluteus medius muscle rate of energy expenditure is plotted throughout the gait cycle for both natural (orange) and exotendon (purple) runs. (b) The peak muscle activation (ranging from 0–1), fiber velocity (fiber lengths per second), and active fiber force averaged across all participants at the time of highest energy expenditure for both natural (orange) and exotendon (purple) running.

## IV. Discussion

The purpose of this study was to investigate muscle-level changes that enable humans to save energy when running with an exotendon, a spring that connects their legs. Using musculoskeletal simulations, we found that the savings from an exotendon during running occurred primarily in the stance phase and resulted from savings in the quadriceps, hip flexor, hip abductor, hamstring, hip adductor, and hip extensor muscle groups. By storing and returning elastic energy during the gait cycle, the spring lowers the rate of energy expenditure required to run at the same speed. Runners took advantage of the mechanical work done by the spring by adopting a gait pattern that reduced both the mechanical work done and the heat generated by muscles.

Our simulation framework accurately captured between-condition changes in whole-body rate of energy expenditure at the group and individual levels that were measured using indirect calorimetry. Further, between-condition changes in simulated muscle activations generally matched those measured with EMG. This provided confidence that the simulations could accurately elucidate muscle-level changes in energy expenditure, providing insights into the human-device interactions that resulted in energy savings. The kinematic and kinetic changes we observed matched those reported by Simpson and colleagues [13] as well. This framework is publicly available, enabling others to explore the muscle-level mechanisms underlying how humans leverage assistive devices.

Despite the exotendon applying force primarily during the swing phase, participants predominantly saved energy during the stance phase. Furthermore, the muscle group that saved the most energy—the quadriceps muscles—did so when the exotendon was not applying any force on the body. These savings were facilitated by the kinematic and kinetic changes adopted when running with the exotendon, including a decrease in knee range of motion and knee extension moment. The smaller knee range of motion required a smaller fiber velocity in the quadriceps muscles such as the vastus lateralis. The reduced knee extension moment also led to a reduction in the force required by the vastus muscle fibers. These changes combined to reduce the required rate of mechanical work done by the vastus lateralis, resulting in energetic savings. Further, the reduction in fiber velocity and activation combined to reduce the rate that heat was generated in the muscle, which additionally saved energy.

Although the exotendon elicited smaller energetic savings during the swing phase, compared to the stance phase, some muscles did save energy during swing. The hip flexors saved energy during swing due to the forces applied by the exotendon. As Simpson et al. [13] showed, the exotendon undergoes two peaks in tension during a single gait cycle. These occur when the legs are at the maximum distance from one another, just before heel-strike and just after toe-off. After toe-off, the hip flexors were aided in accelerating the leg forward from the tension in the exotendon. At the end of swing, just before heel-strike, the hamstrings were aided in accelerating the leg backward due to the tension pulling posteriorly on the leg. In the hip flexors specifically, the reduction in force, due to a reduced hip flexion moment, and fiber velocity, due to a reduced hip flexion range of motion, combined to reduce muscle work. Further, the reduction in fiber velocity and activation combined to reduce the heat generated by the muscle.

The savings in the hip abductors were similar in nature to the knee extensors and hip flexors. With the exotendon, runners had a smaller adduction-abduction range of motion and biological abduction moment. These changes led to a reduction in the fiber velocity and force, which reduced the work rate in the abductors. Further, the reduced fiber velocity and activation produced less heat in these muscles, leading to additional energy expenditure savings. Similar savings mechanisms can be found in other muscles in the model.

It is important to acknowledge the limitations of our simulation approach and analysis. In this work, we used an accepted model of energy expenditure [34], [35]. However, Koelewijn & van den Bogert [38] showed that the model’s ability to accurately capture changes in the magnitude of full-body rate of energy expenditure was limited. Additionally, we did not include muscles in the torso or upper extremities, which likely impacts the estimates of full-body rate of energy expenditure. We believe the results are still valuable, as the simulated kinematics closely match those measured in experiments, the energy costs from simulation were close to measured values, and the simulations correctly detected between-condition changes in rate of energy expenditure. Our muscle-level conclusions are based on changes in energy rate, so matching the raw energy rates is less important than accurately capturing changes.

We have made all the data and results from this study available for download and further analysis at (https://simtk.org/projects/exotendon_sims). Future work should explore how this device assists gait at different speeds and in different configurations. The current and other configurations of a simple exotendon could provide alternative benefits such as offloading a particular joint or tendon. Our simulation framework is generally applicable for studying the effect of devices on gait and could even be adapted to predict gait adaptations to a device [32]. Future studies can leverage this framework to efficiently design devices and device parameters that optimize movement objectives, like energy economy.

## V. Conclusion

We used musculoskeletal modeling and optimal control methods to simulate the effects of a spring attached to the feet that improves energy economy during running. These simulations enabled us to investigate the muscle-level changes that underlie the whole-body reduction in rate of energy expenditure when using the device. We hypothesized that savings would occur in both stance and swing, and that savings would occur in the hip flexors, hamstrings, and quadriceps. We found that while running with the device, runners saved more energy during stance (though savings in swing did occur), and that the savings occurred in the quadriceps, hip flexor, hip abductor, hamstrings, hip adductor, and hip extensor muscle groups. The simulations allowed us to understand the muscle state changes that contributed to the reductions in rate of energy expenditure. This approach can be used to study human-device interactions more generally and could be extended to design novel devices.

## Acknowledgments

We thank Julie Muccini, Nick Bianco, and Cara Welker for their assistance.

